# The effect of intrauterine growth restriction on the developing pancreatic immune system

**DOI:** 10.1101/2024.09.19.613902

**Authors:** Thea N. Golden, James P. Garifallou, Colin C. Conine, Rebecca A. Simmons

## Abstract

Immune cells in the pancreas are known to participate in organ development. However, the resident pancreatic immune system has yet to be fully defined. Immune cells also play a role in pathology and are implicated in diseases such as diabetes induced by intrauterine growth restriction (IUGR). We hypothesized that the resident immune system is established during neonatal development and disrupted by IUGR. Using single cell RNAseq and flow cytometry we identified many immune cell populations in the near-term fetus (at embryologic day 22) and neonatal (postnatal day 1, 7, &14) islets, non-endocrine pancreas, and the spleen in the rat. Using flow cytometry, we observed the resident immune system is established during neonatal development in the pancreas and spleen. We identified 9 distinct immune populations in the pancreatic islets and 8 distinct immune populations in the spleen by single cell RNAseq. There were no sex-specific differences in the relative proportion of immune cells in the pancreas or spleen. Finally, we tested if IUGR disrupted the neonatal immune system using bilateral uterine artery ligation. We found significant changes to the percentage of CD11B+ HIS48- and CD8+ T cells in the islets and non-endocrine pancreas and in the spleen. IUGR-induced alterations were influenced by the tissue environment and the sex of the offspring. Future research to define the role of these immune cells in pancreatic development may identify disrupted pathways that contribute to the development of diabetes following IUGR.

## INTRODUCTION

Previous studies suggest that pancreatic development is dependent on resident macrophages; however, little is known about their function and activity during development. Banaei-Bouchard, et al. first reported the role of macrophages in pancreatic development. Using a macrophage deficient mouse (colony stimulated factor-1 knockout (op/op)), they observed that islets were smaller but more numerous, and pancreatic duct proliferation was increased in macrophage deficient mice^1^. This result suggests that a loss of macrophages result in arrested β-cell proliferation and increased islet neogenesis during development.

The resident immune system in the pancreas is complex and the tissue microenvironment is a key determinant of immune cell phenotype and activity. There are both innate and adaptive immune cells in normal healthy pancreatic tissue ^2,3^. In the mouse, resident immune cells populate the pancreas during mid gestation on embryologic day 14.5 (e14.5) ^4^. Immune cells are initially derived from a yolk-sac precursor and then from a hematopoietic precursor ^5^. After initial establishment during fetal development, resident macrophage turnover is predominately from bone marrow-derived precursors, but evidence exists that macrophages in the exocrine compartment can repopulate from a pancreatic precursor ^5^. Macrophages acquire their phenotype and activation state during neonatal development. In the mouse, resident macrophage MHCII expression increases over the first 4 weeks of life ^6^. Similarly, in the rat islet, the immune system shifts from Th2 driven pathways to Th1 driven pathways from e19 to postnatal day 14 ^7^. In adult mice, macrophages in the endocrine compartment are classically activated (M1) and express pro-inflammatory proteins and high levels of CD64, CX3CR1, CD11c, and MHCII ^5,6^. Macrophages in the exocrine compartment are alternatively activated (M2) and express anti-inflammatory and pro-resolution proteins and lower levels of CD11c and MHCII ^5,6^. Interestingly, lymphocytes are also found in the pancreas, but very little is known about their homeostatic function. Whitesell, et al. recently determined that there are many subsets of lymphocytes, and that the dominant population expresses IL-10 in the neonatal period ^8^.

However, questions remain regarding the role of resident immune cells in tissue homeostasis, development and disease states. It is well established that intrauterine growth restriction (IUGR) causes ß-cell failure leading to the eventual development of T2D. We have developed a model of IUGR that leads to the development of T2D in adulthood. Using this model, we and others have determined that IUGR causes systemic and pancreatic inflammation early in life which is key to the development of β-cell failure ^7,9–14^. However, immune cell populations altered by IUGR, are unknown.

Here we sought to determine the effect of IUGR on the normal establishment of resident immune cells. We found that pancreatic immune cell composition differed between the endocrine and non-endocrine compartments in normal developing pancreas. We also observed that IUGR altered these endocrine vs. non-endocrine compartment differences in early life in a sex-specific manner.

## MATERIALS AND METHODS

### Pancreas and Spleen Sample Collection

The animal care committees of the Children’s Hospital of Philadelphia and University of Pennsylvania approved all animal use and procedures. Pregnant Sprague-Dawley dams were purchased from Charles River for all animal experiments. All euthanasia was performed by intraperitoneal injection of 100mg/kg ketamine and 10mg/kg xylazine.

Each litter was a biological replicate, and males and females were analyzed separately to test the role of sex in normal development and following intrauterine growth restriction (IUGR). Each biological replicate was generated by pooling tissues from offspring of the same sex from the same litter to ensure sufficient tissue for measurement. The pancreas and spleen were analyzed from the same rats. At the late fetal stage, embryologic day 22 (e22), pregnant dams were injected with ketamine and xylazine and pups were excised from the uterus. Sex was determined by anogenital distance, and the pancreas and spleens were excised. To ensure sufficient sample for measurement at e22, all tissues from either males or females of the same litter were combined to generate a male and female sample from each litter. To assess offspring post birth, pregnant rats were allowed to spontaneously deliver the litter. Pups were injected with 100mg/kg ketamine and 10mg/kg xylazine followed by decapitation. At postnatal day 1 (PD1), tissues were excised from three pups of the same sex and pooled to generate a male and female sample from each litter. At PD7, the tissues of two same sex offspring were pooled to generate a male and female sample from each litter. Finally, the tissue of a single rat at PD14 was sufficient for measurement which eliminated the need to pool. Each litter was treated as a biological replicate and three litters were analyzed at e22, PD1, and PD7 and six litters were analyzed at PD14.

### Intrauterine Growth Restriction Animal Model

To measure the effect of intrauterine growth restriction on the resident immune system in the pancreas and spleen, we used the bilateral uterine artery ligation method. Pregnant Sprague Dawley rats underwent bilateral uterine artery ligation at day 18 of gestation, as previously described ^7,15,16^. In brief, pregnant rats were anesthetized with isoflurane and pain was managed with local bupivacaine (2mg/kg SQ) and tramadol (12.5mg/kg IP) before surgery and postoperatively. The uterine artery was isolated, and silk suture was used to tie the artery between the first amniotic sac and the uterine horn, bilaterally. Rats recovered from surgery and spontaneously delivered pups. Pups born to dams that underwent surgery (IUGR) or no surgery (controls) were weighed on postnatal day 0 and litters were culled to 7-9 pups per litter to achieve uniformity. IUGR resulted in a 35.3% decrease in male and 40.9% decrease in female birth weight (males: control 8.5g ± 0.6g compared to IUGR 5.5g ± 1.1g and females: controls 8.3g± 0.6g compared to IUGR 4.9g ± 1.3g). Dams were given ad libitum access to standard rat chow and water.

### Immune Cell Isolation from Pancreas and Spleen

Fetal pancreata were digested with 0.3mg/mL Collagenase XI (Sigma C7657) and PD1-PD14 pancreas were digested with 0.6mg/mL in DMEM with 5% FBS at 37°C for 12 minutes with intermittent shaking. Tissue was spun down at 500G for 2 minutes and pellet resuspended in HBSS. Cell suspension was filtered through a 425um mesh filter (Bellco Glass 1985-00040) and washed with HBSS. Cell pellet was resuspended in Histopaque 1.077 (fetal: 1.2mL and PD1-PD14: 3.4mL) and Histopaque 1.119 (fetal: 2.8mL and PD1-PD14: 7.5mL) was added to the suspension. Histopaque 1.119 (fetal: 4mL and PD1-PD14: 12mL) was layered under and Histopaque 1.077 was layered on top ((fetal: 2.8mL and PD1-PD14: 7.5mL). Finally, 4.2uM NaHCO3 1% BSA HBSS (fetal: 4mL and PD1-PD14: 12mL) was layered on top. The gradient suspension was spun at 855G with no brake at 4°C for 25 minutes. Islets were visualized in the interphase and aspirated. The pellet was collected and washed in HBSS (non-endocrine portion of pancreas).

Intact islets were further purified by adding 20mL 4.2uM NaHCO3 1% BSA HBSS and spinning at 500G at 4°C for 2 minutes twice. The pellet was resuspended in 1mL Cell Dissociation Solution (Sigma C5789) which was prewarmed to 37°C and 1mg/mL DNAse I-Grade II (Roche 10104159001) by pipetting the cell suspension ten times. Cells were incubated at 37°C for 5 minutes. The resulting cell suspension were washed in PBS and the pellet collected (islets).

Spleens were excised from rats and cell suspensions were generated using a plunger (BD 309585). The resulting cell suspension was pipetted up and down in DMEM to further break apart the tissue before filtering through a 40um filter (Falcon 352340). Cells were then spun at 450G at room temperature for 5 minutes. The resulting pellet was resuspended and incubated with ACK lysis buffer (Gibco A1049201) for 1 minute at room temperature to lyse red blood cells. PBS was added to the tube to stop lysis and spun at 450G at 4°C for 5 minutes and the pellet resuspended in PBS.

### Immune Cell Isolation and staining for Flow Cytometry

Immune cells were isolated from cell suspensions using the StemCell EasySep magnetic assisted cell sorting. Cells were incubated with FcR block (Mouse Anti-Rat Cd32 BDBiosciences 550270, RRID:AB_393567) at 50uL/mL followed by biotinylated CD45 antibody (BD Biosciences 554876, RRID:AB_395569) at 1ug/mL. Cd45+ cells were isolated following manufacturer EasySep Biotin Positive Selection Kit (StemCell 17655) protocol and the EasyPlate EasySep magnetic plate (StemCell 18102). Resuspended cells were manually counted using a hemacytometer.

### Flow Cytometry

Cells were resuspended at 1×10^7^/100uL and co-incubated with fluorescent antibodies following manufacturer protocol. Antibodies include CD3 (BD Biosciences 563949, RRID:AB_2738504), CD45Ra (BD Biosciences 740726, RRID:2740404), CD4 (BD Biosciences 740256, RRID:AB_2740000), CD8 (BD Biosciences 740041, RRID:AB_2739811), CD11B (Biolegend 201809, RRID:313995), HIS48 (BD Biosciences 743057, RRID:AB_2741252). BD Fortessa II flow cytometer was used to measure fluorescence and FlowJo software for analysis. Single cells were identified by SSC-H and SSC-A gating and live cells based on Live/Dead Blue (Invitrogen L23105) staining. The complete gating strategy is shown in Supplementary Figure 1. Isotype controls were used as negative controls.

### Immune Cell Isolation and staining for Flow Assisted Cell Sorting

Pancreatic islets from PD7 rats were digested as described above. 2 rats were used to generate each sample and 3 samples were prepared. Immune cells were isolated from single cell suspensions via Miltenyi Biotec Magnetic Cells Assisted Sorting. Cells were resuspended in 80uL buffer and 20uL CD45 microbeads (Miltenyi Biotec 130-109-682) at 4°C for 15 minutes before adding to a prepared Pre-Separation Filter (130-101-812) and MS column (130-042-201). Cells were collected after filtration and spun down at 400G at 4°C for 5 minutes.

Cells were resuspended at 1×107/100uL and co-incubated with fluorescent antibodies following manufacturer protocol. Antibodies include CD3 (BD Biosciences 550353, RRID:AB_393632), CD45Ra (BD Biosciences 740726, RRID:2740404), CD11B (BD Biosciences 562222, RRID:AB_11154584), HIS48 (BD Biosciences 554907, RRID:AB_395595) at 4°C for 30 minutes. Cells were washed and incubated with 1uL Live/Dead Violet dye (Invitrogen L23105) 4°C for 20 minutes. Cells were washed and resuspended in 50% FBS in stain buffer. Cells were sorted on the FACSAria II to collect live CD45+ CD3- CD45Ra- CD11B+ HIS48+ and CD45+ CD3- CD45Ra- CD11B+ HIS48+ single cells in 100% FBS. Cells were spun at 500G at 4°C for 5 minutes and resuspended in RNAlater ICE (Invitrogen AM7030).

### Bulk RNASeq in Flow-sorted cells

To further characterize the immune cell population in normal animals during development, we used a separate cohort of animals. RNA was isolated from flow-sorted cells with Qiagen RNAeasy kit (Qiagen 74104) following manufacturer protocol. mRNA libraries were prepared from total RNA using the SMART-Seq protocol ^17^. In brief, RNA was isolated via RNAClean-XP (Beckman Coulter, Cat#A63987) and full length polyadenylated RNA was reverse transcribed using Superscript II (Invitrogen, Cat#18064014). cDNA was amplified with 10 cycles and amplified with the Qubit dsDNA HS Assay Kit (Life Technologies, Inc. Cat#Q32851). 0.33ng of each sample was used to construct a pool of uniquely identified samples with the Nextera XT kit (Illumina Cat# FC-131-1096) and a second amplification of the pooled samples was conducted with 12 cycles. The pooled sample was cleaned up with AMPure XP beads and the final library was sequenced on a NextSeq 1000. Data was mapped against the rat_rn6_refseq genome using DolphinNext pipeline and a STAR ^18^.

To quantitate and normalize the expression data and then assess the differentially expressed genes (DEGs) the data was loaded into R statistical software and analyzed using DESeq2. When CD11B+ HIS48+ to CD11B+ HIS48- cells expression, genes with a log2 fold change greater than 1.5 and an adjusted p-value less than 0.05 were considered significant.

### Immune cell isolation and staining for single cell RNAseq preparation

PD1 animals were euthanized by decapitation and pancreas and spleens were isolated as described above. Cells were isolated from 5 males and 5 females as described above. Cells were pooled to generate 2 pancreas and 2 spleen samples from both males and females. Cells were resuspended in 50uL BV buffer and 100uL buffer and 1uL FcR block was added and incubated at 4°C for 5 minutes. CD45 antibody (BD Biosciences 561586, RRID:AB_10896305) was added and incubated at 4°C for 30 minutes. Cells were washed in PBS and spun down at 500G at 4°C for 5 minutes. Cells were resuspended in PBS and 1uL Live/Dead Violet dye (Invitrogen L23105) for at 4°C for 20 minutes. Cells were washed in PBS and spun down at 500G at 4°C for 5 minutes. Cells were resuspended in buffer.

Cells were sorted by the FACSAria II and CD45+ Live cells were collected in 10% FBS in DMEM. 28,000-63,000 cells were collected for each pancreas sample and 100,000 cells were collected for each spleen sample. Cells were spun at 500G at 4°C for 5 minutes and resuspended in PBS. Viability was then confirmed to be above 85% and cells were counted.

### Single Cell RNASeq Library Preparation

Next generation sequencing libraries were prepared at The Center for Applied Genomics at Children’s Hospital of Philadelphia using the 10X Genomics Chromium Single Cell 3’ Library & Gel Bead Kit v2 as per manufacturer instructions (PN-120237). cDNA and library preparation quality were confirmed with the Agilent Bioanalyzer High Sensitivity Kit and final concentration was determined with a KAPA Library Quantification Kit (PN-07960140001). Libraries were individually indexed, pooled, and sequenced on the Illumina Hiseq2500 using SBS chemistry v4 in a paired-end, single indexing run.

### Single Cell RNASeq Data processing and Analysis

Data was demultiplexed and processed using the cellranger mkfastq and count pipelines (10x genomics, v.6.1.2) for demultiplexing, alignment of sequencing reads to the rattus norvegicus transcriptome (mRatBN7.2), and creation of feature-barcode matrices. Secondary analysis was performed using the Seurat package (v.5.0) ^19^ within the R computing environment. Spleen and pancreas datasets were analyzed separately. Each dataset was filtered for cell quality to include cells with greater than or equal to 250 UMIs, 250 genes expressed, and mitochondrial content less than 20 percent. Data was merged, normalized, and the top 2000 variable features were used for principal component analysis. The harmony package in R ^20^ was used on the PCA to account for per library batch effects in downstream UMAP and clustering (resolution 0.5). Small clusters expressing high levels of collagen and red blood cell genes were determined to be contaminating clusters and were removed from dataset. Cell types were annotated using canonical marker genes. The dittoSeq R package was used for additional figures ^21^.

### Statistical analysis

Statistical analysis was performed via GraphPad Prism version 10. Flow cytometric data was tested for normality with the Shapiro-Wilk test. Data that passed normality was tested by a one-way ANOVA followed post hoc pairwise analysis with Fisher’s test. Data that failed normality was tested by Kruskal-Wallis test followed by post hoc analysis with Dunn’s test. Values that exceeded 2x the standard deviation were considered outliers and removed from analysis. P-values less than 0.05 were considered significant and the p-values for significant tests were recorded on the figure above the comparison groups. Gene expression was considered significant if the log2 fold change exceeded 1.5 and the adjusted p-value below 0.05.

## RESULTS

### The immune cell composition in normal pancreas during development

The pancreatic resident immune system is populated during early life. We analyzed samples in the near-term fetus (e22), and neonate (PD1, PD7, and PD14) as these end-points capture the window of time in which the immune system shifts from a predominantly Th2 to a Th1 phenotype. In adults, distinct immune populations exist in the endocrine and exocrine pancreas, thus we enumerated immune cells in the endocrine (islets) and exocrine (non-endocrine portion) pancreas separately. We used a panel of antibodies to detect the major classes of immune cells including B cells (CD45ra+), T cells (CD3+), and myeloid cells (CD11B+) (Supplemental Figure 1). To further delineate the T cell population, we identified the expression of CD4 and CD8 on CD3+ cells. Myeloid cells were further demarcated based on their expression of HIS48.

In islets, the dominant immune cell population was CD11B+ HIS48- (Figure 1A) at all time points. This population gradually decreased over the first two weeks of life. The CD11B+ HIS48+ population was stable until a significant decrease at PD14 (Figure 1B). The percentage of both CD4+ and CD8+ T cells was very low at e22 and PD1 but gradually increased over time, whereas B cell numbers fluctuated significantly at each time point (Figure 1 C-E). Interestingly, there was no effect of sex on the relative percentage of immune cells.

**Figure 1.**
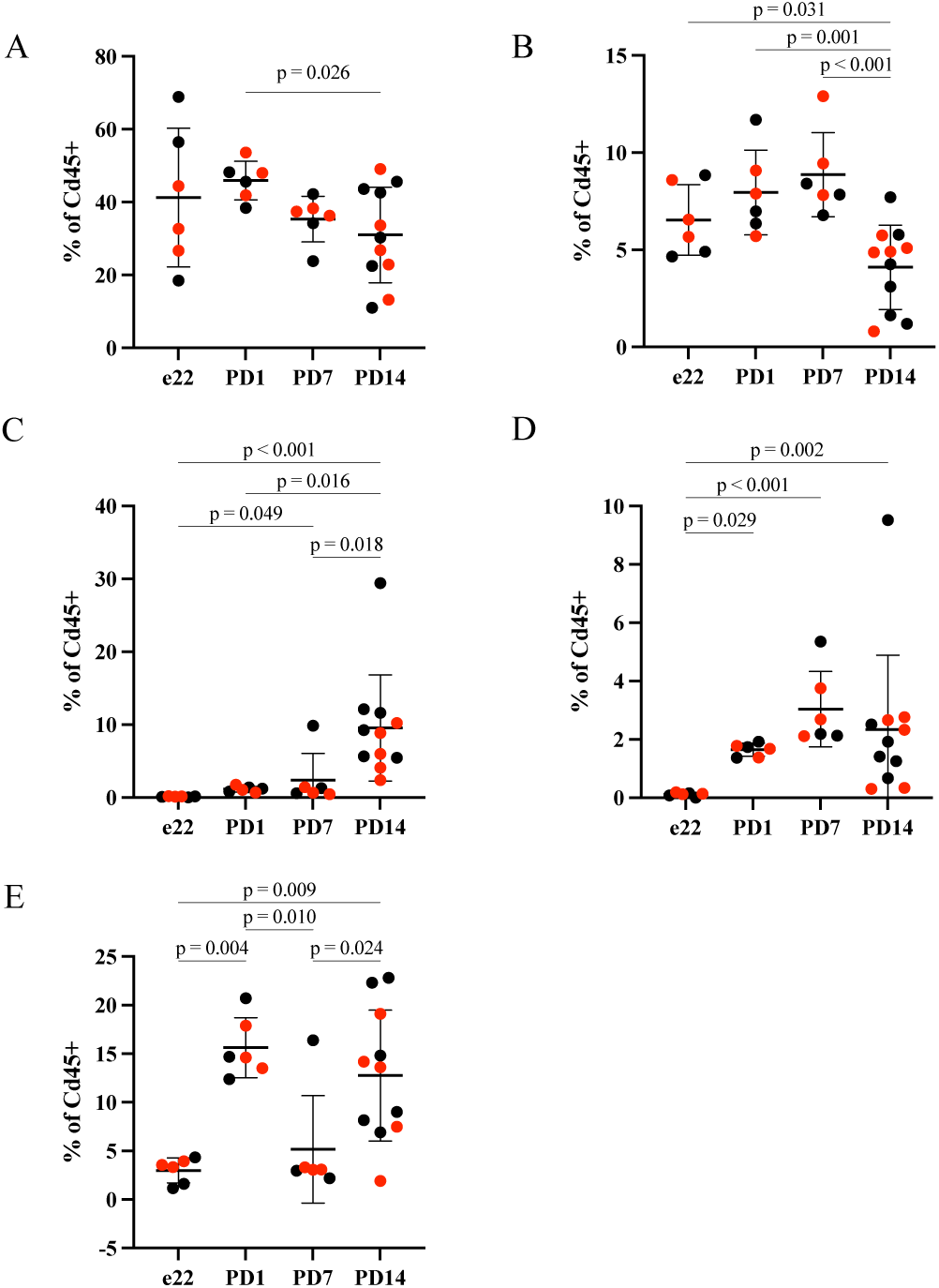
Immune cells were isolated from pancreatic islets were identified by protein expression. The relative proportion of immune cells in the pancreatic islet are reported as the percentage of CD45+ cells: CD11B+HIS48- (A), CD11B+HIS48+ (B), CD4+ T cells (C), CD8+ T cells (D), B cells (E) in males (black) and females (red). Comparison groups are identified by a bar and the p-value recorded for those with a significant post hoc test.

Many immune cells were also abundant in the non-endocrine portion of the pancreas and the dominant population was CD11B+ HIS48- (Figure 2A). The percentage of CD11B+ HIS48+ cells decreased over time and was lowest at PD14. The percentage of CD11B+ HIS48+ cells gradually increased with age and plateaued at PD7 (Figure 2B). T cells were very low in abundance at e22 and PD1 but expanded with age (Figure 2C&D). B cells in the non-endocrine pancreas shared a similar pattern of fluctuation over neonatal life to those in the islets (Figure 2E). There was also no sex-specific effect on the percentage of immune cells in the non-endocrine portion of the pancreas.

**Figure 2.**
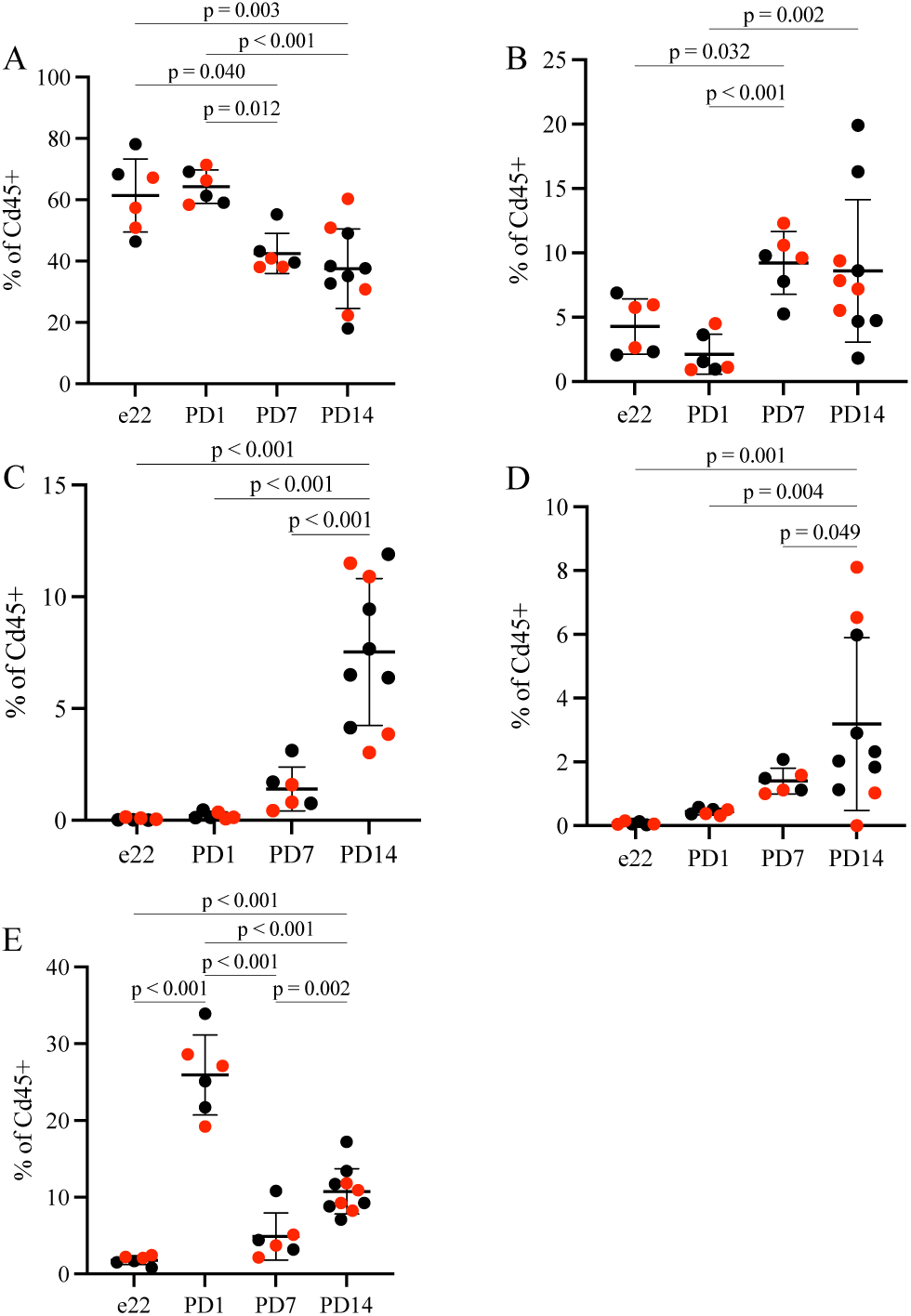
Immune cells isolated from the non-endocrine pancreas were identified by flow cytometric detection of surface proteins. The relative proportion of immune cells in the non-endocrine pancreas are reported as the percentage of CD45+ cells: CD11B+HIS48- (A), CD11B+HIS48+ (B), CD4+ T cells (C), CD8+ T cells (D), B cells (E) in males (black) and females (red). Comparison groups are identified by a bar and the p-value recorded for those with a significant post hoc test.

The CD11B+ cell population clearly separated into HIS48- and HIS48+ subpopulations in islets and non-endocrine portion of the pancreas. While HIS48 expression is commonly used in rats to identify myeloid cells, its biological function is unknown. In other rat tissues, HIS48 expression identified subsets of macrophages and granulocytes ^22,23^. To identify the phenotype of myeloid cells based on HIS48 expression we analyzed the transcriptome of flow sorted CD11B+ HIS48- and CD11B+ HIS48+ cells. There were 26 differentially expressed genes (DEG) between CD11B+ HIS48+ compared to CD11B+ HIS48- cells; 17 genes had higher expression and 9 had lower expression (Figure 3). The genes with increased expression in CD11B+ HIS48+ cells encode for pro-inflammatory proteins that are known to be highly expressed in neutrophils and classically activated monocytes/macrophages including, *CD177, Plyrp1, Camp, S100A8, S100A9, Olm4, Chit1, Hp, Padj4,* and *Fcnb* (Table 1). Surprisingly, CD11B+ HIS48- cells had high expression of genes commonly expressed by mast cells including *Tpsb2, Tpsab1, Mcpt2, Cpa3, Mcpt1l1,* and *Cma1* (Table 1). The transcriptome of the two subpopulations suggests the population that expressed CD11B+ HIS48+ contained pro-inflammatory myeloid cells and the population that expressed CD11B+ HIS48- cells contained mast cells.

**Figure 3.**
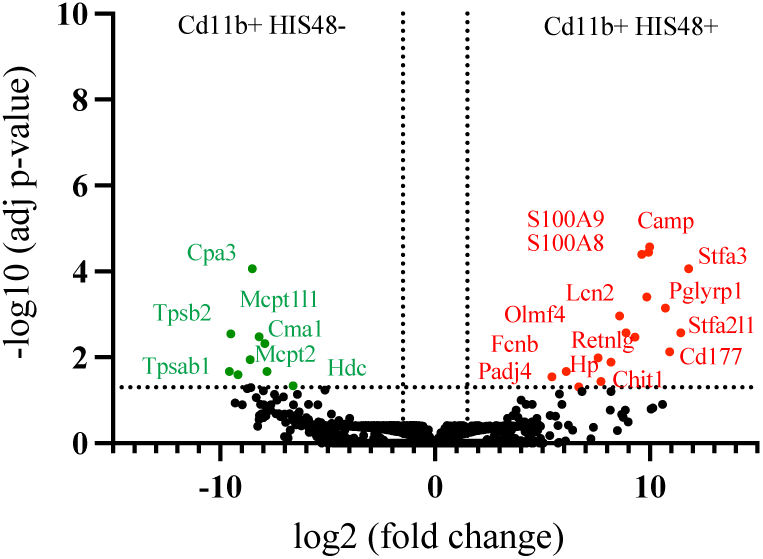
Comparing the gene expression of CD11B+ HIS48+ to CD11B+ HIS48- flow sorted cells. Genes with expression greater than log2 fold change and an adjusted p-value less than 0.05 are colored red. Genes with expression less than log2 fold change and an adjusted p-value less than 0.05 are colored green.

**Table 1.**
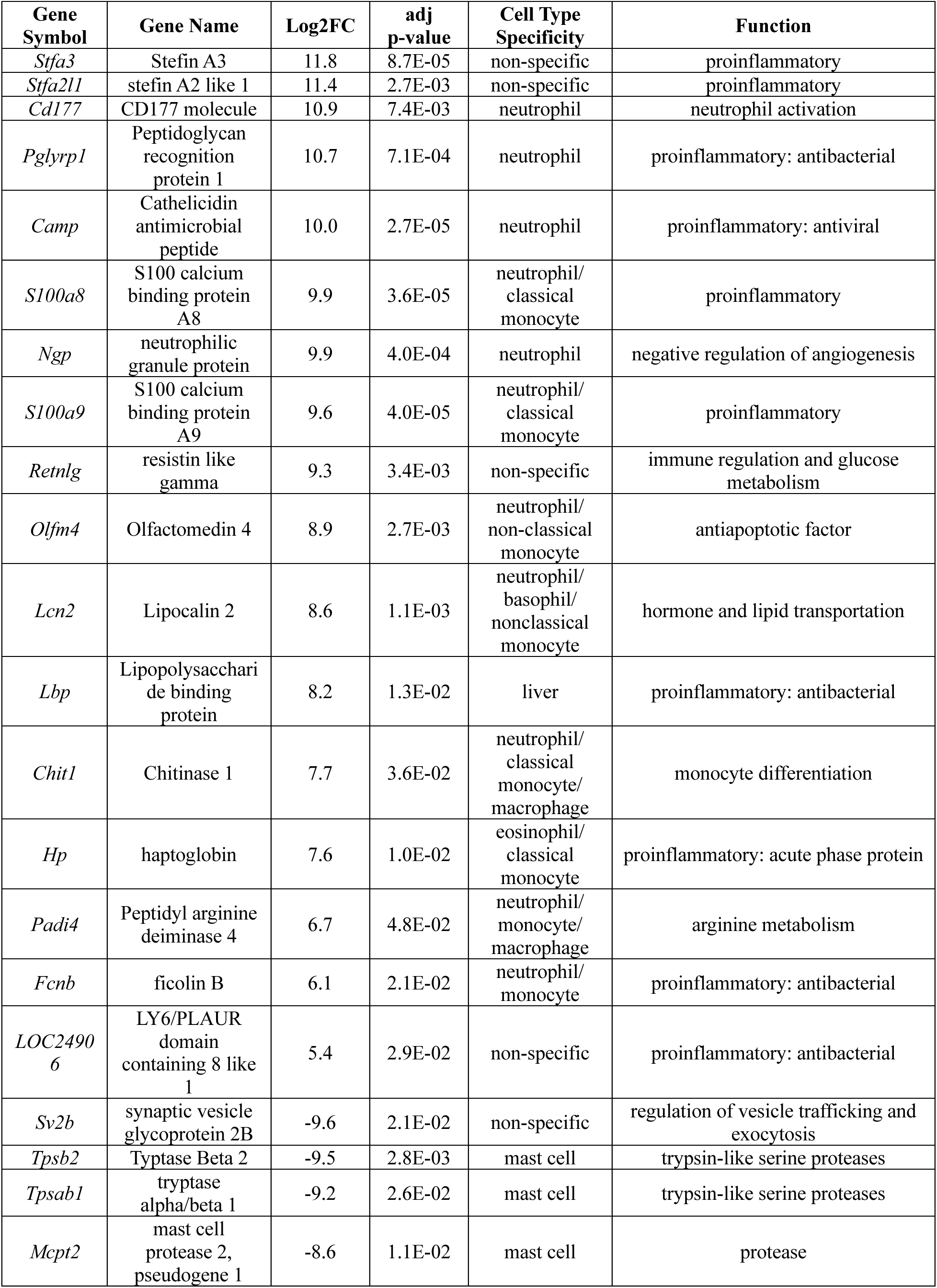
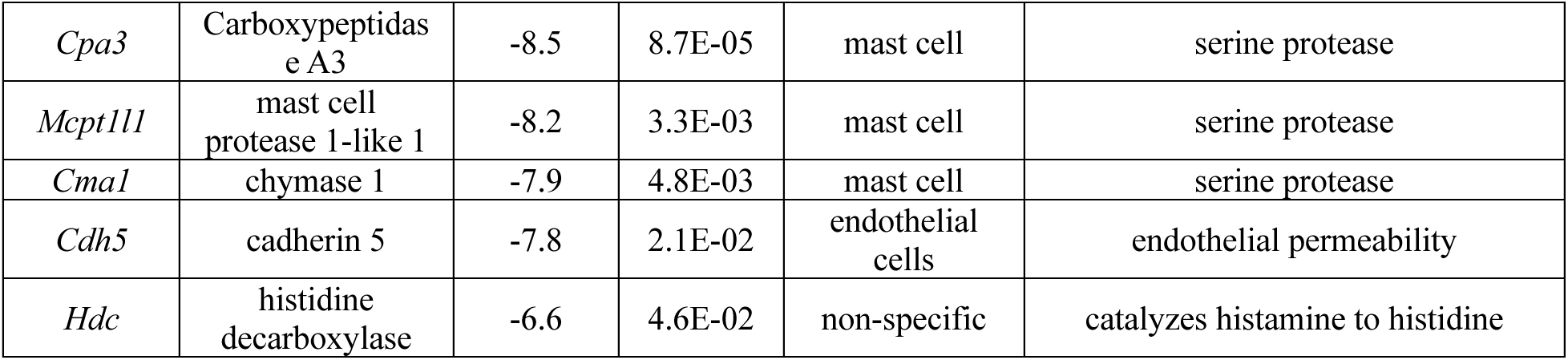
Differentially expressed genes were identified by RNAseq comparing CD11B+HIS48+ compared to CD11B+HIS48- cells. The log 2-fold change, adjusted p-value are listed for each gene. The specific immune cell type with high expression and the functional role of each gene is listed.

| Gene Symbol | Gene Name | Log2FC | adj p-value | Cell Type Specificity | Function |
| --- | --- | --- | --- | --- | --- |
| *Stfa3* | Stefin A3 | 11.8 | 8.7E-05 | non-specific | proinflammatory |
| *Stfa2l1* | stefin A2 like 1 | 11.4 | 2.7E-03 | non-specific | proinflammatory |
| *Cd177* | CD177 molecule | 10.9 | 7.4E-03 | neutrophil | neutrophil activation |
| *Pglyrp1* | Peptidoglycan recognition protein 1 | 10.7 | 7.1E-04 | neutrophil | proinflammatory: antibacterial |
| *Camp* | Cathelicidin antimicrobial peptide | 10.0 | 2.7E-05 | neutrophil | proinflammatory: antiviral |
| *S100a8* | S100 calcium binding protein A8 | 9.9 | 3.6E-05 | neutrophil/<br>classical monocyte | proinflammatory |
| *Ngp* | neutrophilic granule protein | 9.9 | 4.0E-04 | neutrophil | negative regulation of angiogenesis |
| *S100a9* | S100 calcium binding protein A9 | 9.6 | 4.0E-05 | neutrophil/<br>classical monocyte | proinflammatory |
| *Retnlg* | resistin like gamma | 9.3 | 3.4E-03 | non-specific | immune regulation and glucose metabolism |
| *Olfm4* | Olfactomedin 4 | 8.9 | 2.7E-03 | neutrophil/<br>non-classical monocyte | antiapoptotic factor |
| *Lcn2* | Lipocalin 2 | 8.6 | 1.1E-03 | neutrophil/<br>basophil/<br>nonclassical monocyte | hormone and lipid transportation |
| *Lbp* | Lipopolysaccharide binding protein | 8.2 | 1.3E-02 | liver | proinflammatory: antibacterial |
| *Chit1* | Chitinase 1 | 7.7 | 3.6E-02 | neutrophil/<br>classical monocyte/<br>macrophage | monocyte differentiation |
| *Hp* | haptoglobin | 7.6 | 1.0E-02 | eosinophil/<br>classical monocyte | proinflammatory: acute phase protein |
| *Padi4* | Peptidyl arginine deiminase 4 | 6.7 | 4.8E-02 | neutrophil/<br>monocyte/<br>macrophage | arginine metabolism |
| *Fcnb* | ficolin B | 6.1 | 2.1E-02 | neutrophil/<br>monocyte | proinflammatory: antibacterial |
| *LOC24906* | LY6/PLAUR domain containing 8 like 1 | 5.4 | 2.9E-02 | non-specific | proinflammatory: antibacterial |
| *Sv2b* | synaptic vesicle glycoprotein 2B | -9.6 | 2.1E-02 | non-specific | regulation of vesicle trafficking and exocytosis |
| *Tpsb2* | Typtase Beta 2 | -9.5 | 2.8E-03 | mast cell | trypsin-like serine proteases |
| *Tpsab1* | tryptase alpha/beta 1 | -9.2 | 2.6E-02 | mast cell | trypsin-like serine proteases |
| *Mcpt2* | mast cell protease 2, pseudogene 1 | -8.6 | 1.1E-02 | mast cell | protease |
| *Cpa3* | Carboxypeptidase A3 | -8.5 | 8.7E-05 | mast cell | serine protease |
| *Mcpt1l1* | mast cell protease 1-like 1 | -8.2 | 3.3E-03 | mast cell | serine protease |
| *Cma1* | chymase 1 | -7.9 | 4.8E-03 | mast cell | serine protease |
| *Cdh5* | cadherin 5 | -7.8 | 2.1E-02 | endothelial cells | endothelial permeability |
| *Hdc* | histidine decarboxylase | -6.6 | 4.6E-02 | non-specific | catalyzes histamine to histidine |

### The pancreatic immune landscape in the neonate

Our data suggests there are many immune cell populations in the pancreas during early life. To identify subpopulations not captured by our flow cytometry panel, we used single-cell RNAseq to interrogate the transcriptome of immune cells isolated from PD1 rat islets. These immune cells are likely to participate in islet establishment. Single-cell RNA libraries were prepared from immune cells isolated from PD1 islets by flow sorting of CD45+ live cells.

UMAP clustering determined there were 9 distinct immune subpopulations that grouped into the following subclusters: B cell, dendritic cells (DC), macrophages, myeloid cells (5), and natural killer cells (NK) (Figure 4A). Not surprisingly, we did not detect any clusters that expressed high levels of typical T cell markers (*Cd3, Cd4, or Cd8*) as this population was less than 2% of the total immune population quantified by flow cytometry at PD1. B cells expressed high levels of *Cd24, Jchain, Cd79b, Mki67,* and *Tifa* (Figure 1B). There were many clusters that expressed myeloid cell specific markers including *Ftl1, Fcer1g,* and *Lyz2* at varying levels. A clear macrophage population was identified by high expression of these genes and the expression of mannose receptor (*Mrc1)*, a marker of pro-resolution macrophages. Myeloid cluster 1, 2, and 5 also expressed *C1qc, Ftl1, Fcer1g,* and *Lyz2*. Interestingly, myeloid cluster 1 also expressed *Ly6C* which is commonly expressed in pro-inflammatory macrophages. Myeloid cluster 5 also expresses high levels of the DEGs with increased expression in the CD11B+ HIS48+ cells as determined by bulk RNAseq (Figure 4D). The DC cluster expressed *Cd74* and *Cst3* which are highly expressed in dendritic cells. Myeloid cluster 3 also expressed moderate *Cd74* and *Cst3* but also expressed *JChain* suggesting these cells may be plasmacytoid DCs. Myeloid cluster 4 did not express any transcripts that clearly defined its cell type, but interestingly cells that expressed high levels of CD11B+ HIS48- DEGs, as determined by bulk RNAseq, were found in this cluster (Figure 4E). Finally, a clear cluster of natural killer cells were defined by the expression of *Nkg7, Kird1,* and *Ccl5*. The relative percentage of cells in each cluster did not vary by sex, in our limited dataset (Figure 1C).

**Figure 4.**
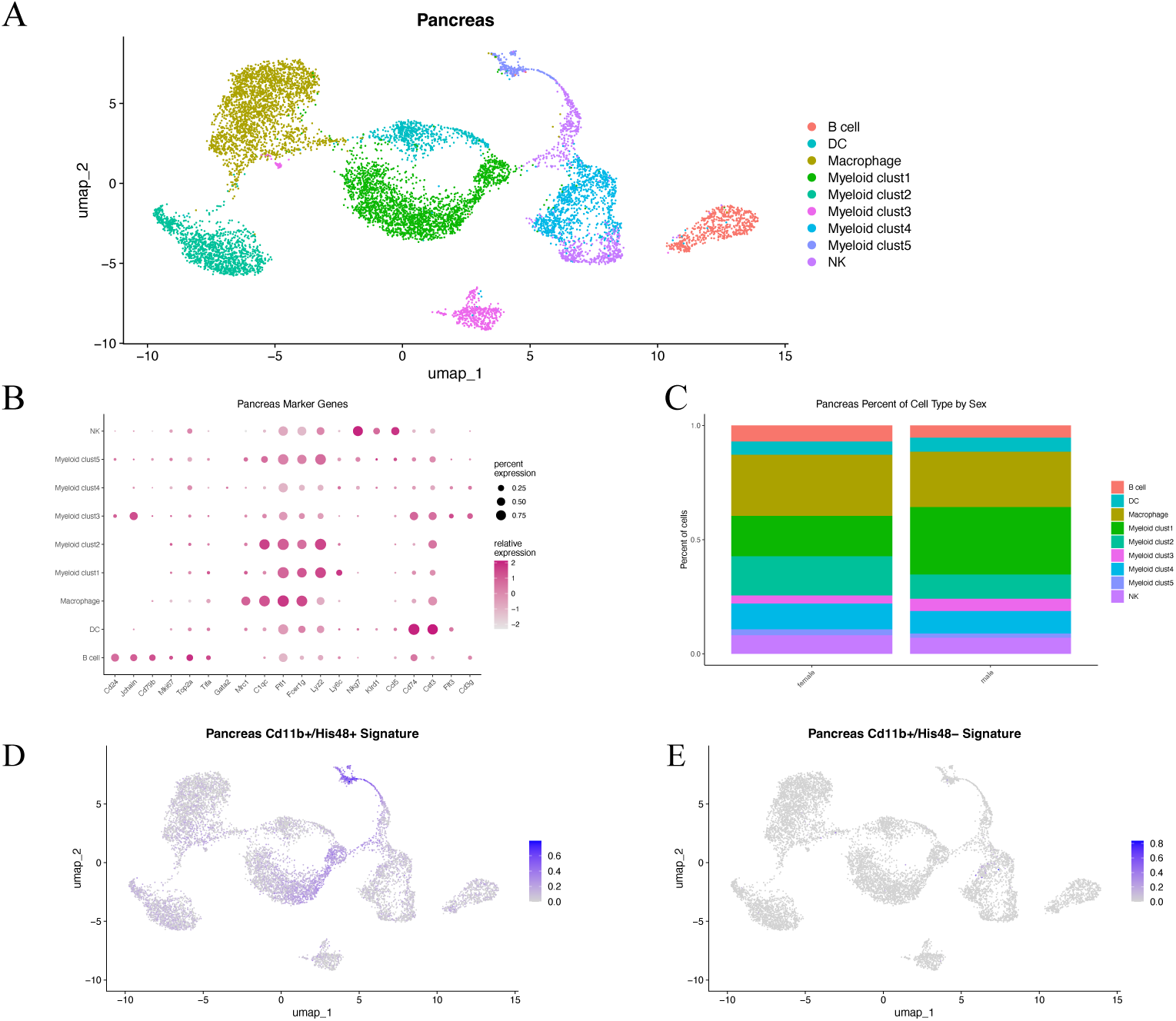
Pancreatic islet immune landscape at postnatal day 1. Pancreatic islets were isolated from pancreata excised from postnatal day 1 male and female neonatal rats. Immune cells were flow sorted based on CD45 expression and negative live/dead dye. Single cell RNA sequencing of flow sorted cells resulted in the identification of 9 distinct clusters (A). The marker genes that drive clustering are listed (B). The percentage of cells in each cluster separated by sex (C). Expression of differentially expressed genes, determined by bulk RNA sequencing of flow sorted CD11B+ HIS48+ (D) and CD11B+ HIS48- (E) are identified in the single cell RNAseq feature plots.

### Immune cells in the neonatal spleen

The spleen is a secondary lymphoid organ that has significant roles in immunity and blood homeostasis. The resident immune system in the murine neonate was recently found to vary considerably from the adult ^24^. Mundim Porto-Pedrosa, et al. observed a gradual decline in B cells but increase in T cells in the neonatal period before the adult immune system was established. The authors also reported the number of macrophages and monocytes transiently increased in the neonatal period but returned to post birth levels in adulthood. To elucidate the establishment of the splenic immune system in the rat, we analyzed immune cells isolated from the spleen at e22, PD1, PD7, and PD14.

At late gestation, the dominant immune cell population in the spleen was CD11B+ HIS48- cells (Figure 5B). Following birth, there was a significant decrease in the percentage of CD11B+ HIS48- cells that increased at PD7. Unlike in the pancreas, the number of CD11B+ HIS48+ cells was very low at birth and further declined with age (Figure 5A). Like in the mouse, B cells are abundant in the spleen, and we found the percentage rapidly increased with age (Figure 5E). Interestingly, in the spleen the percentage of T cells also increased with age, similar to the pancreas (Figure 5C&D). This suggests that, like in the mouse, the resident immune system is not fully established at birth but continues to develop during the neonatal period.

**Figure 5.**
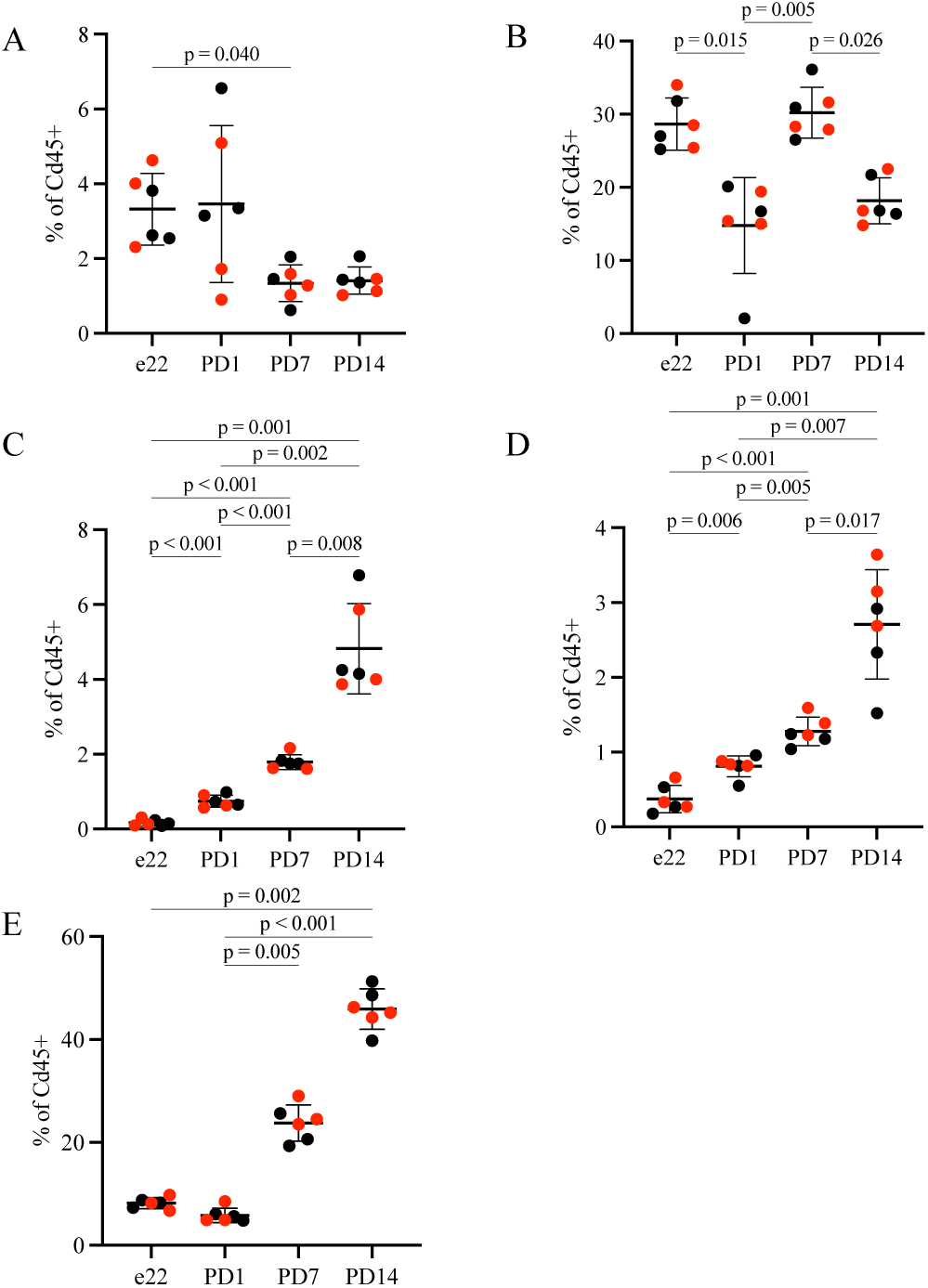
The relative proportion of immune cells in the spleen are reported as percentage of CD45+ cells: CD11B+HIS48- (A), CD11B+HIS48+ (B), CD4+ T cells (C), CD8+ T cells (D), B cells (E) in males (black) and females (red). Comparison groups are identified by a bar and the p-value recorded for those with a significant post hoc test.

### The immune landscape in the neonatal spleen

Many subpopulations of immune cells exist beyond the capacity of our flow cytometry antibody panel to detect. Therefore, we analyzed the transcriptome of immune cells isolated from the spleens at PD1 of the same rats as the pancreas scRNAseq analysis. CD45+ live immune cells were isolated from splenic tissue and single cell RNA libraries were prepared and sequenced. The single cell transcriptome clustered into 8 distinct subpopulations that expressed typical cell identifying genes (Figure 6A). B cells, expressing *Ms4a1, Ly86, Cd24, Cd27, Cd38* and *Cd79b,* comprised over 50% of the immune cells (Figure 2). B cells further subdivided into immature B cells, based on *Vrep3* expression, and proliferating B cells based on *Mki67, Top2a*, and *Tif1* expression (Figure 6B). T cells were a small but identifiable population with expression of *Cd3g*. Expression of myeloid cell type specific genes allowed for clustering into macrophages and neutrophils/monocytes. Macrophages expressed high levels of *C1qc, Ftl1, Fcer1g, and Lyz2.* Cells defined as neutrophils/monocytes expressed high levels of *Ftl1, Fcer1g, Lyz2, and Ly6C*. Interestingly, the neutrophils/monocytes cluster also contained cells that expressed high levels of the DEGs in CD11B+ HIS48+ cells determined by bulk RNAseq (Figure 6D). Gene expression of the DEGs increased in CD11B+ HIS48- cell were found in the cluster termed mast cells (Figure 6E). There was also no sex-related difference to the percentage of cells in each cluster (Figure 2C). It is surprising that more immune cells in the spleen expressed canonical genes that allow for more cell-type identification of clusters than those immune cells in the pancreas. This suggests the tissue microenvironment was a major determinant of gene expression in the neonatal rat.

**Figure 6.**
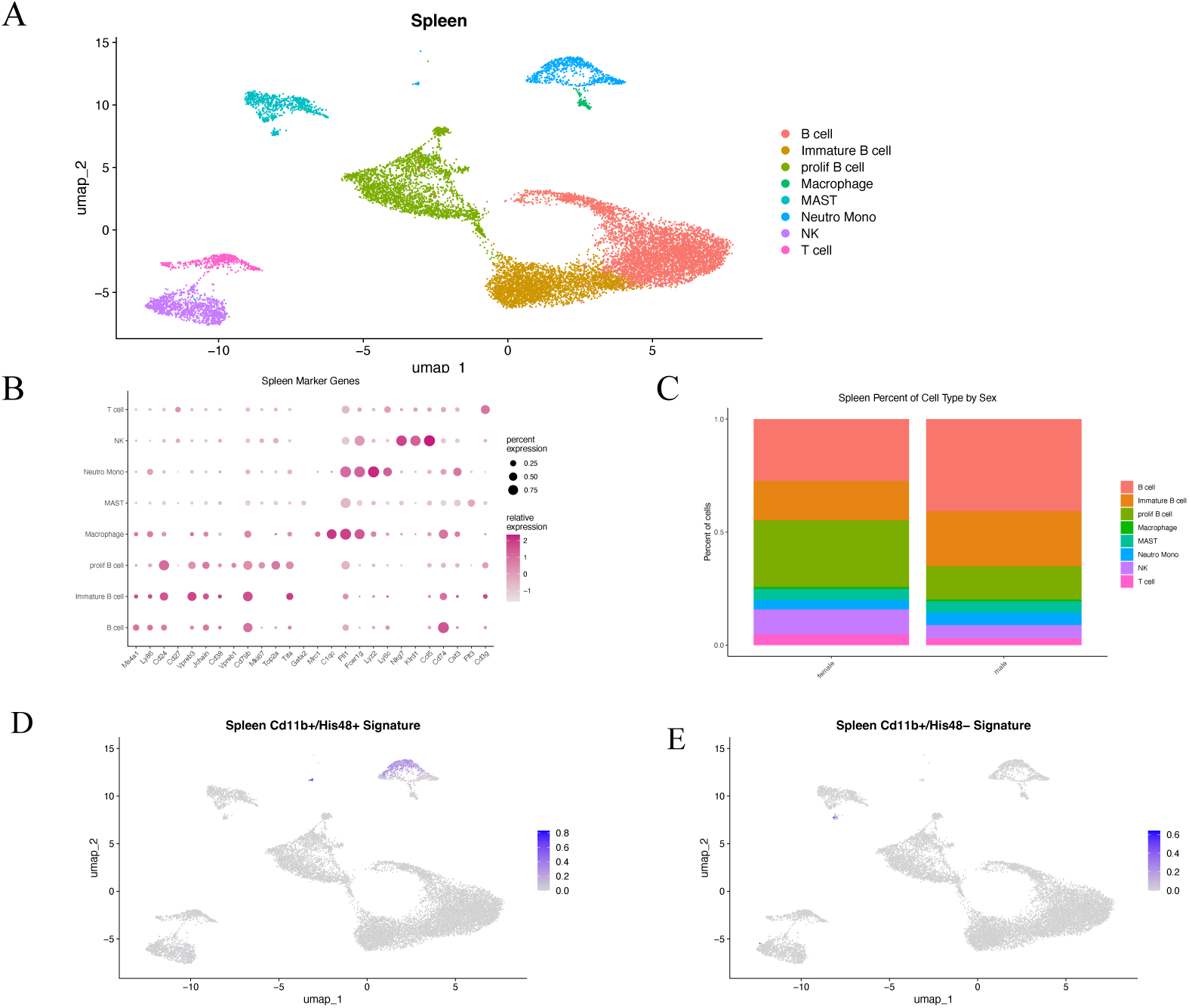
The immune landscape in the spleen at postnatal day 1. 8 total clusters were identified by single cell RNA sequencing of immune cells isolated from the spleen (A). The marker genes that drive clustering are listed (B). The percentage of cells in each cluster separated by sex (C). Expression of differentially expressed genes, determined by bulk RNA sequencing of flow sorted CD11B+ HIS48+ (D) and CD11B+ HIS48- (E) are identified in the single cell RNAseq feature plots.

### Immune cell populations are altered by IUGR

Having observed that there is a dynamic establishment of the pancreatic immune system, we sought to determine if the altered in utero environment elicited by IUGR disrupted normal immune development. The percentage of CD11B+ HIS48- cells at e22 was significantly reduced in islets from IUGR males and females compared to controls (Figure 7A). However, the number of CD11B+ HIS48- cells rapidly increases postnatally and is similar to controls. Simultaneously, there was an increase in the percentage of CD8+ T cells in male and female IUGR offspring at e22. Interestingly, the percentage of CD8+ T cells in females rapidly returned to control levels unlike in the males which took longer (Figure 7A&B). Finally, there was also a transient decrease in the percentage of B cells in the islets isolated from male and female IUGR offspring at PD1 that rapidly returned levels similar to controls by PD7 (Figure 7B&C). These early changes in the relative abundance of immune cells in IUGR islets demonstrate that specific immune cell populations were differentially affected by the altered intrauterine milieu.

**Figure 7.**
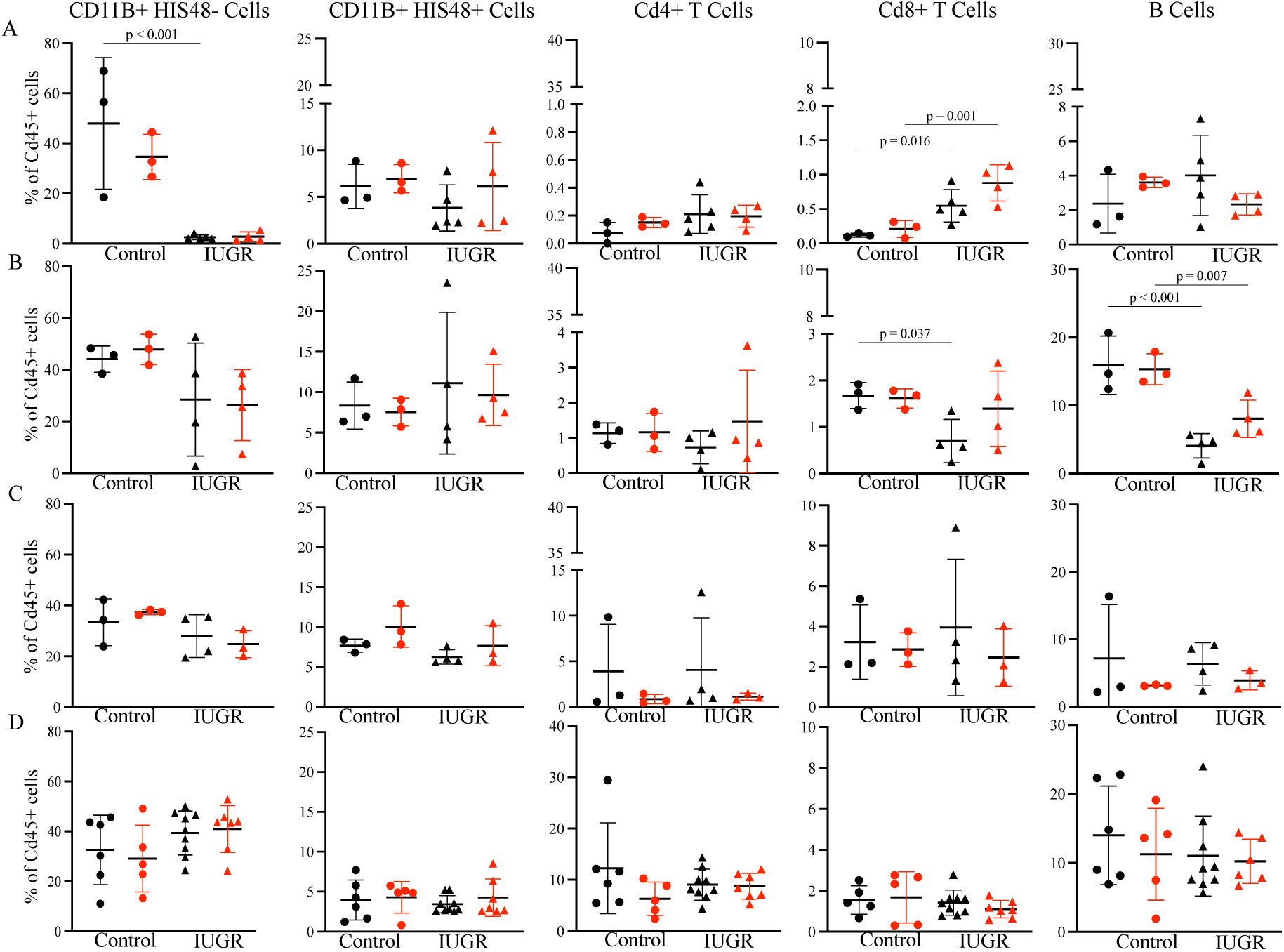
Immune cell composition in the islets following IUGR. Immune cells were quantified at e22 (A), PD1 (B), PD7 (C), and PD14 (D) in males (black) and females (red). Comparison groups are identified by a bar and the p-value recorded for those with a significant post hoc test.

IUGR also had a cell type specific effect on the immune cell composition in the non-endocrine portion of the pancreas. At e22, the percentage of CD11B+ HIS48+ and CD4+ T cells was decreased in female IUGR compared to female control offspring. However, by PD1 the percentage of these populations recovered and did not differ from control female offspring (Figure 8A&B). At PD1, the percentage of CD11B+ HIS48- was increased in male IUGR offspring but interestingly this percentage was decreased at PD7 before reaching a similar percentage to controls at PD14 (Figure 8B-D). Also, at PD1 there was a decrease in the percentage of CD8+ T cells that rebounded at PD7 before normalizing at PD14. IUGR-induced disruption of pancreatic immune composition had a longer lasting effect on the male than female offspring suggesting a sex-specific effect of IUGR. This is particularly important because only males develop glucose intolerance in this model of IUGR.

**Figure 8.**
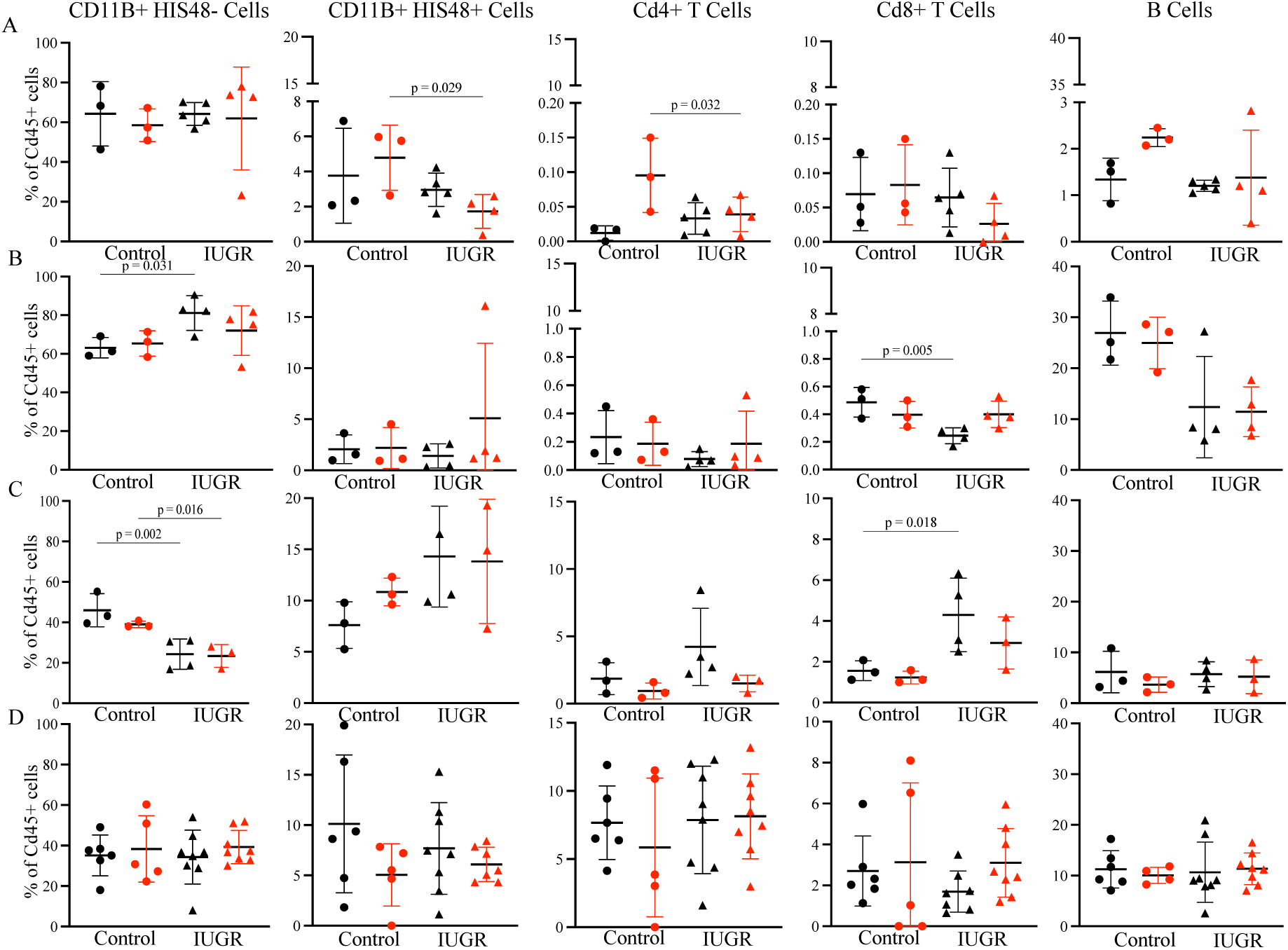
Immune cell composition in the non-endocrine pancreas following IUGR. Immune cells were quantified at e22 (A), PD1 (B), PD7 (C), and PD14 (D) in males (black) and females (red). Comparison groups are identified by a bar and the p-value recorded for those with a significant post hoc test.

Finally, to test if the alterations induced by IUGR were systemic, we analyzed immune cells in the spleen. Interestingly, we saw different changes in the abundance of immune cells in the spleen compared to in the pancreas. CD11B+ HIS48, CD4+ T cells, and CD8+ T cells were reduced in IUGR female offspring at e22 but were similar to controls postnatally (Figure 9A).

**Figure 9.**
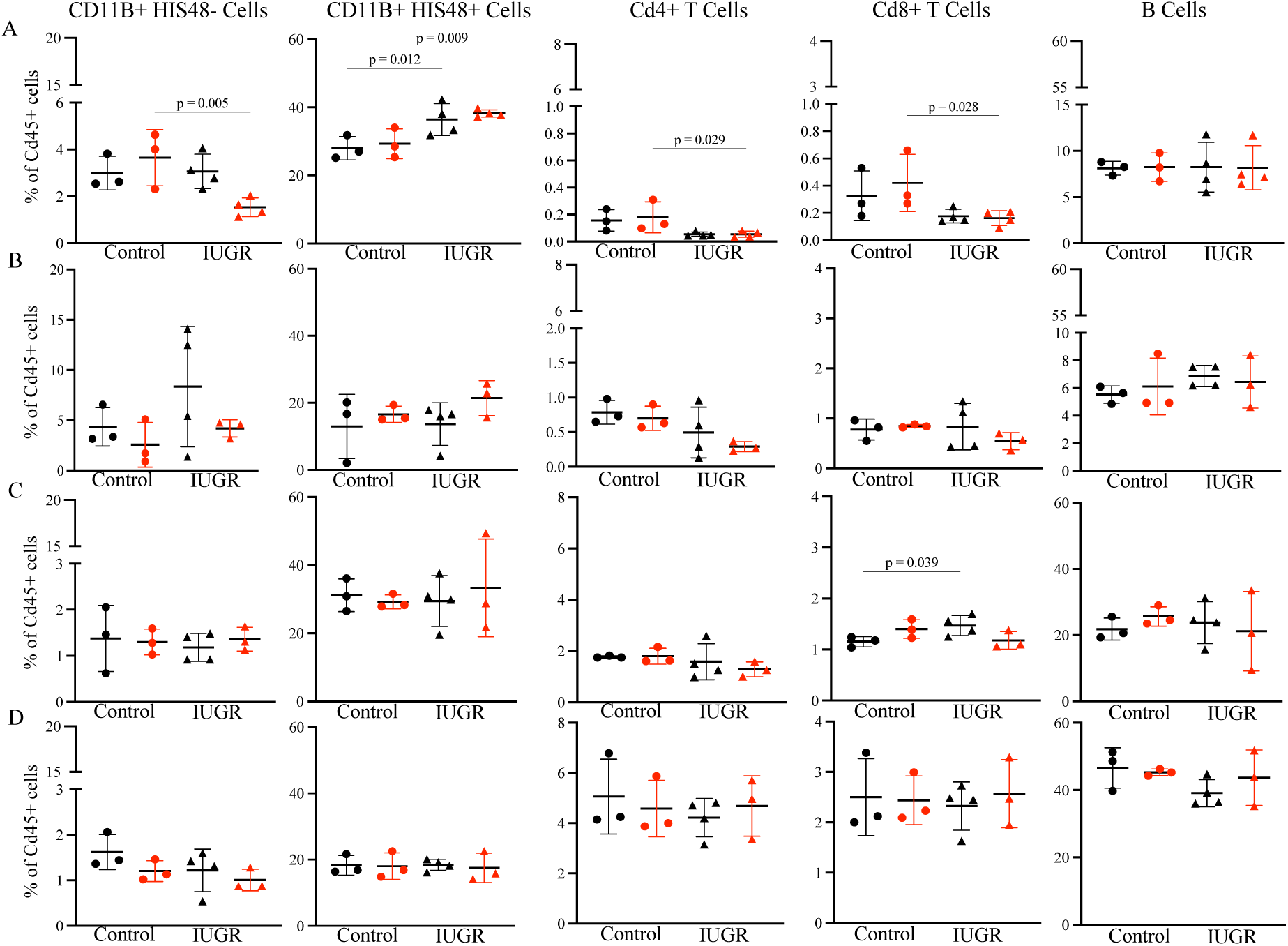
Immune cell composition in the spleen following IUGR. Immune cells were quantified at e22 (A), PD1 (B), PD7 (C), and PD14 (D) in males (black) and females (red). Comparison groups are identified by a bar and the p-value recorded for those with a significant post hoc test.

Only CD11B+ HIS48+ cells were increased in both male and female IUGR offspring at e22 but were similar in number to controls postnatally (Figure 9A&B). Finally, there was a transient increase in the percentage of CD8+ T cells in male but not female IUGR offspring at PD7 (Figure 9C). Like in the pancreas, all populations were normalized by PD14 (Figure 9D). These results demonstrate the influence of the cellular microenvironment on the effect of IUGR.

## DISCUSSION

In this study, we quantified and analyzed the resident immune cell populations in the neonatal pancreas for the first time. We found that the pancreatic immune cell landscape significantly changes during the neonatal period. The dominant population in both islets and non-endocrine pancreas were myeloid cells, but the percentage of T and B cells increased with age. We also confirmed that the resident immune system in the rat continued to develop in the neonatal period as previously reported in the mouse. At birth, the majority of residential immune cells were myeloid, but B cells quickly expanded postnatally. Finally, we found significant sex-specific effects of IUGR on the developing pancreatic immune system but only minimal disruption of the developing splenic immune system.

Myeloid cells, particularly macrophages, are known to play a vital role in organ development ^25^. Macrophages promote tissue remodeling, angiogenesis, and apoptosis induction through cytokine and growth factor release ^26^. Studies using a pancreatitis model in rodents showed an important role of immune cells in tissue remodeling and demonstrate that immune cells participate in pancreatic development ^27,28^. The precise pathways and immune cells vital to pancreatic development remain to be identified ^29^.

Islet remodeling is a normal part of pancreatic development during the first two weeks of life in rodents ^30,31^. During fetal and early neonatal development, new β-cells are predominately formed via differentiation of embryonic ductal cells. In the later postnatal period and into adulthood, new β-cells are formed via replication of preexisting β-cells. The replication rate is low in adulthood but does continue with age. The molecular mediators of β-cell neogenesis and differentiation have been well described. In addition, macrophages have been shown to play a vital role in islet formation. Macrophages participate in pancreatic innervation through phagocytosis of apoptotic nerves ^32^. It has also been demonstrated that macrophages clear apoptotic β-cells which is part of the process of islet remodeling ^33^.

We observed that the majority of immune cells in the neonatal pancreas are myeloid-derived, and many are macrophages. Similar to what has been observed in mice, the percentage of myeloid-derived cells decreased over the neonatal period in rats ^34^. However, the canonical markers used to delineate myeloid subtypes were not highly expressed in the immune cells isolated from the pancreas thus limiting our ability to identify subtypes.

Surprisingly, we found numerous CD11B+ HIS48- cells in the pancreas that express high levels of protease transcripts commonly detected in mast cells. During early pancreatic development, mast cell recruitment to the pancreas may be responsive to pancreatic epithelial expression of a potent mast cell chemoattractant, transforming growth factor beta (TGFβ) ^35^. Upon recruitment, mast cells release mediators such as tumor necrosis factor alpha (TNF-ɑ), histamine, metalloproteinase 9 (MMP9), and interleukin 4 (IL4)—all of which participate in organ development and angiogenesis in other tissues ^36^. We detected these mediators in myeloid cluster 5 in the single cell RNAseq dataset and found that they are differentially expressed by sorted CD11B+ HIS48- cells compared to other myeloid derived cells (CD11B+ HIS48+). Moreover, in the presence of acute pancreatic inflammation, mast cells participate in the regeneration of pancreatic ducts ^37^. Together these previous studies and our findings suggest that mast cells play an important role in pancreatic tissue remodeling.

IUGR disrupts immune cell composition in the offspring pancreas. In both male and female offspring, CD11B+ HIS48- cells are reduced at e22 in islets. In contrast, IUGR had an inverse effect on CD8+ T cells and increased the number of CD8+ T cells at e22. Moreover, in the non-endocrine pancreas in IUGR males, the percentage of CD11B+ HIS48- cells was initially increased at PD1 but reduced at PD7. At these same time points, the percentage of CD8+ T cells were initially reduced and then increased. The bi-directional changes in the percentage of CD11B+ HIS48- and CD8+ T cells at the same time points suggest that there may be an interaction between these cell types. The transcriptome of CD11B+ HIS48- cells demonstrated this population contains mast cells. Interestingly, mast cells and T cells release mediators that induce migration and activation of each other ^38,39^.

Finally, we observed that IUGR only acutely altered the percentage of immune cells in the spleen. At embryologic day 22, the percentage of mast cells, proinflammatory myeloid cells, CD4 T cells, and CD8 T cells were altered in female IUGR offspring, but the percentage was similar to controls postnatally. This early response may reflect systemic changes to the immune system following IUGR and highlights the susceptibility of the pancreas to the lasting effects of IUGR.

In conclusion, we observed a complex pancreatic resident immune system that is unique from the spleen. Subpopulation percentages changed over the neonatal period, suggesting an active role of the immune system during this critical window of pancreatic development. IUGR disrupted immune cells in both male and female offspring, but the rate of recovery was slower in male offspring. Future research focused on understanding the role of these immune cells, particularly CD11B+ HIS48- and CD8 T cells, in pancreatic development may identify novel pathways responsible for islet failure following IUGR.

## ACKNOWLEDGEMENTS

The authors would like to thank Dr. G. Scott Worthen for his invaluable contribution to the study design and interpretation of results.

## Funding

National Institute of Health grant R01DK114054 (RAS)

National Institute of Health grant T32ES019851 (TNG)

National Institute of Health grant P30ES013508 (TNG)

## Author contributions

Experimental Design: TNG, RAS

Data acquisition and analysis: TNG, JPG, CCC

Writing - original draft: TNG, RAS

Writing - review & editing: TNG, JPC, CCC, RAS

## Competing interests

The authors declare they have no competing interests.

## DATA AVAILABILITY

### Data and materials availability

The sequencing data reported in this study are deposited in GEO. All other data are available in the main text.

---

Supplemental Figure 1. Immune cell types were identified by cellular surface protein expression. Single cells were gated to only include live CD45+ cells. T cells were identified by CD3 expression. CD3+ cells were further analyzed to quantify CD4 and CD8 T cells. CD3- cells were further gated on CD45Ra to identify B cells. CD3- CD45Ra- cells were further gated on CD11B and HIS48 to quantify myeloid cells.

## Notes

### Competing Interest Statement

The authors have declared no competing interest.

