## Supplementary figures and images for "The effect of intrauterine growth restriction on the developing pancreatic immune system"

### Supplemental Figure 1

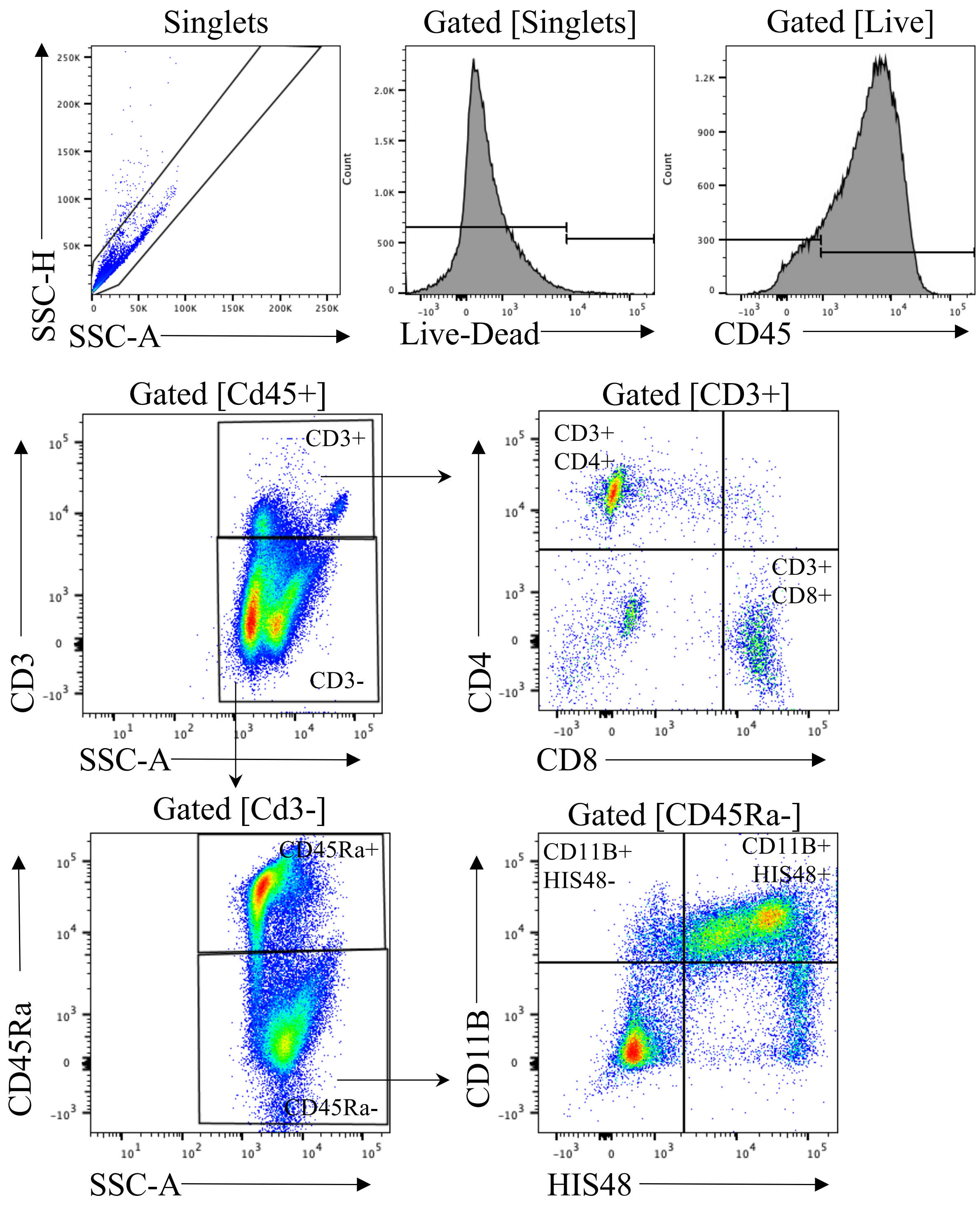
